# Probabilistic representations as building blocks for higher-level vision

**DOI:** 10.1101/2021.11.18.469104

**Authors:** Andrey Chetverikov, Árni Kristjánsson

## Abstract

Current theories of perception suggest that the brain represents features of the world as probability distributions, but can such uncertain foundations provide the basis for everyday vision? Perceiving objects and scenes requires knowing not just how features (e.g., colors) are distributed but also where they are and which other features they are combined with. Using a Bayesian computational model, we recover probabilistic representations used by human observers to search for odd stimuli among distractors. Importantly, we found that the brain integrates information between feature dimensions and spatial locations, leading to more precise representations compared to when information integration is not possible. We also uncover representational asymmetries and biases, showing their spatial organization and arguing against simplified “summary statistics” accounts. Our results confirm that probabilistically encoded visual features are bound with other features and to particular locations, proving how probabilistic representations can be a foundation for higher-level vision.

## Introduction

How the brain represents the visual world is a long-standing question in cognitive science. One captivating idea is that the brain builds statistical models that describe probability distributions of visual features in the environment (Rao et al., 2002; Pouget et al., 2000; Zemel et al., 1998; R. D. Lange et al., 2020; Fiser et al., 2010; Knill & Pouget, 2004; Tanrıkulu et al., 2021). By combining information about different features and their locations, the brain can then form representations of objects and scenes. Indeed, the idea that the brain represents feature distributions matches our conscious visual experience well. Most objects, such as the apple in Figure 1A, contain a multitude of feature values that can be quantified as a probability distribution, and we are seemingly aware of these feature constellations. Surprisingly, most studies of probabilistic representations do not test how such constellations are represented, assuming instead that a stimulus is described by a single value, such as the orientation of a Gabor patch in vision studies or the hue of an item in working memory experiments and that the only uncertainty comes from the sensory noise. While this unrealistic assumption was noted (Zemel et al., 1998) early on, it is still prevalent, leaving open the possibility that the results can be explained with alternative models without assuming detailed representations of probability distributions (Block, 2018; Cohen et al., 2016; Rahnev, 2017).

**Figure 1.**
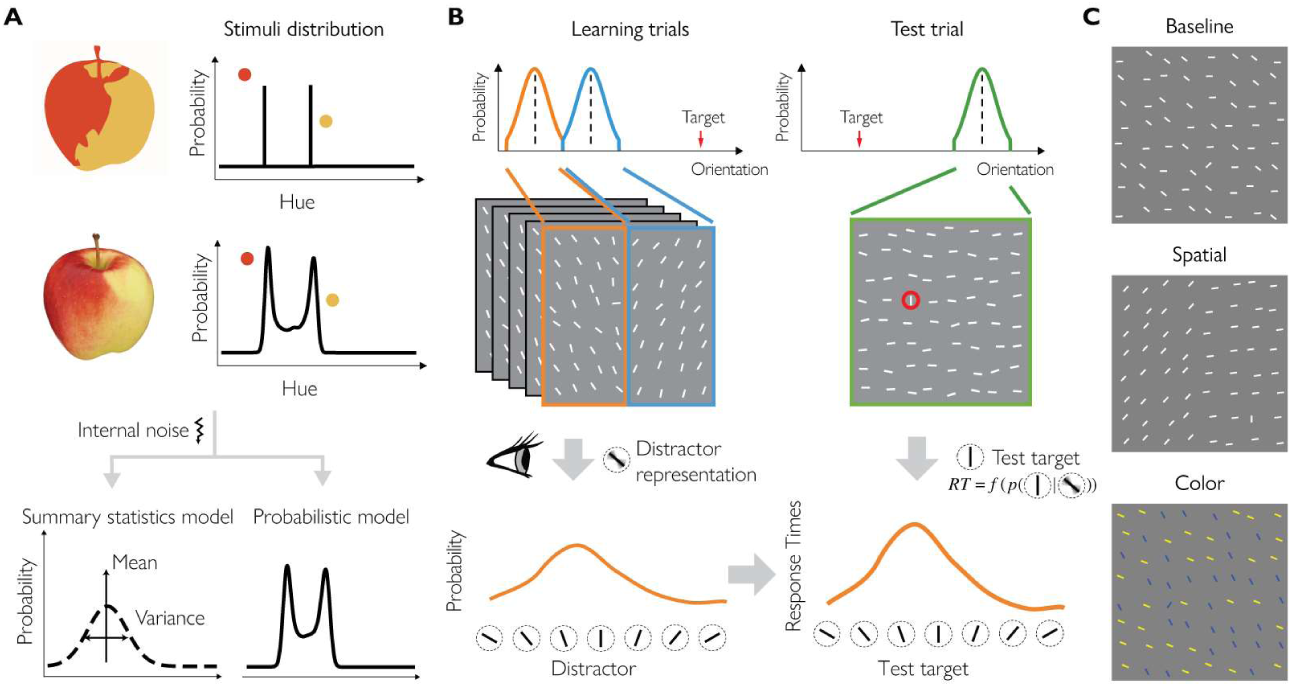
General approach and methods. A: A typical stimulus used to study probabilistic perception involves an impoverished version of the environment, similar to a sketch of an apple (top-left). The hues of this stimulus can be quantified as a discrete probability distribution with only a few probable values (top-right). In contrast, real objects have a multitude of feature values corresponding to a complex-shaped probability distribution (middle). An accurate probabilistic model would maintain the important details of the distribution as much as internal noise permits, while a summary statistics model suggests that probabilities are represented as a combination of simple parameters, such as mean and variance (bottom). B: In our experiments, in each block observers searched for an odd-one-out line among distractors. On learning trials (upper-left), distractors were drawn from two distributions that were either mixed together or separated by location or color with one example of the spatial separation shown here. We assumed that observers would form a distractor representation by learning which distractors are more probable (bottom-left). On test trials (upper-right), we varied the similarity between the target and previously learned distractors. We then measured response times assuming that they should be monotonically related to the probability of a given target being a distractor based on a simplified ideal observer model (bottom-right). C: Example stimuli used on learning trials in Experiment 1.

Here, we aim to close this gap and ask 1) if the visual system is capable of quickly forming precise representations of heterogeneous stimuli, representations that reflect the probability distribution of their features and 2) if such representations can be bound to other features or to spatial locations thereby serving as building blocks for upstream object and scene processing.

How can the brain represent heterogeneous stimuli, that is, stimuli that have more than one feature value? The visual system may track each feature value at each location to form a representation that would be identical to the stimulus. However, this would be extremely costly in terms of computational resources and unnecessary or even misleading for action because specific feature values can vary from one moment to another because of changes in viewpoint, lighting, etc. Another possibility is that only a few values, for example, the mean and the variance (“summary statistics” Ariely, 2001; Cohen et al., 2016; Haberman & Whitney, 2012; Rahnev, 2017; Treisman, 2006), are represented. But this is also unlikely because multiple stimuli can have the same summary statistics while being quite different from each other. More realistically, the brain could follow the middle course by approximating feature distributions in the responses of neuronal populations that capture the important aspects of stimuli without being too detailed (Figure 1A).

Previous studies have indeed shown that the visual system encodes the approximate distribution of visual features and uses them in perceptual decision-making (Girshick et al., 2011; Seriès & Seitz, 2013). However, most of the findings are confined to relatively long-term learning of environmental statistics. If feature probability distributions are to be useful for everyday visual tasks, such as object recognition or scene segmentation, the brain needs to learn feature distributions quickly and effortlessly. Importantly, we have previously provided initial evidence that such rapid learning may occur in simple cases by studying how human observers learn to ignore distracting stimuli while searching the visual scene (Chetverikov et al., 2016, 2017b, 2019, 2020). Observers were asked to find an odd-one-out item in a search array where distractor features (colors or orientations) were randomly drawn from a given probability distribution for several trials. A test trial was then presented with a target of varying similarity to previously learned distractors. We found that response times as a function of this similarity parameter followed the shape of the previously learned probability distribution, whether it was Gaussian, uniform, skewed, or even bimodal. That is, the search was slowed proportionally to how unexpected the target was, based on previously learned environmental statistics. This shows that representations of the shape of feature probability distributions in the visual input (similar to scene statistics(Oliva & Torralba, 2001; Rosenholtz, 2016)) is not limited to long-term learning, but can occur rapidly.

This previous work was, however, limited to simple scenarios with a single feature distribution present, while real environments contain multiple objects and scene parts with different features. Furthermore, knowledge about statistics of a given feature (e.g., orientation) in isolation is not very useful. Observers need to know *where* in the external world a given feature distribution is and which other features should be bound with it (related to the “binding” problem, Treisman, 1996) to recognize objects or segment scenes. Notably, such binding to spatiotopic locations and to other features does not necessarily require any additional neural machinery, because information about feature distributions can be readily encoded in neural population responses (Pouget et al., 2000; Sahani & Dayan, 2003; Vértes & Sahani, 2018; Zemel et al., 1998). Evidence for such effortless integration of probabilistic visual inputs is, however, still lacking.

Ensemble averaging studies testing how observers estimate probabilistic properties of several sets of stimuli provide some initial support for this hypothesis. It is well known that observers can estimate the average of a perceptual ensemble, such as the mean orientation of a set of lines (Alvarez, 2011; Haberman & Whitney, 2012; Whitney & Yamanashi Leib, 2018). Furthermore, they can estimate properties of subsets grouped by location or by other features although this causes performance detriments (Attarha et al., 2014; Attarha & Moore, 2015a, 2015b; Chong & Treisman, 2005; Oriet & Brand, 2013; Utochkin & Vostrikov, 2017). However, this approach has only provided evidence for single-point estimates (the mean) but not for representations of feature probability distributions. Here, we aim to fill this gap and test how observers encode properties of feature distributions and associate them with both spatial locations and other features.

## Results

In three experiments, observers viewed dressed-down versions of the environment that allowed precise control of the critical aspects of feature distributions. Observers searched for an unknown oddball target that differed from other items in orientation and judged whether it was in the upper or lower half of the stimulus matrix (Figure 1B). Observers did this quickly and accurately despite not knowing the target or distractor parameters in advance (average response time across experiments and conditions *M* = 754 ms, *SD* = 197, proportion correct *M* = 0.90, *SD* = 0.04).

In all experiments, trials were organized in blocks of intertwined learning and test trials. In each block, during five to seven learning trials distractor stimuli were drawn randomly from the same probability distribution. On test trials, we varied the similarity of the current target to non-targets from preceding trials (Figure 1B). Using this data, we aimed to understand how observers represent complex heterogeneous stimuli such as visual search distractors.

### Bayesian observer model

How do behavioral responses depend on distractor representations from previous trials? To answer this question and to reconstruct distractor representations from the behavioral responses of our observers, we built a Bayesian memory-guided observer model linking observers’ internal representations of distractors to response times.

Our participants located a target among a set of distractors and indicated if it is in the top or the lower part of the stimuli matrix. On each trial, the experimenter sets the parameters of the target feature distribution, *p*(*s*_*i*_|*L*_*T*_ = *i*), and of the distractor feature distribution, *p*(*s*_*i*_|*L*_*T*_ ≠ *i*), for each location *i* = 1 … *N* in the stimuli matrix as well as the target location (*L*_*T*_). These parameters are then used to generate the stimuli at each location (*s*_*i*_). Neither the task parameters nor the stimuli are known to the observer.

Instead, at each moment in time *t*, the observer obtains sensory observations at each location (*x*_*i,t*_). These observations are not identical to the stimuli because of sensory noise, *p*(*x*_*i,t*_|*s*_*i*_). In other words, a given stimulus might result in different sensory responses, and, conversely, a given sensory observation might correspond to different stimuli.

To find the target, the observer compares for each location the probability that the sensory observations are caused by a target present at that location, *p*(*L*_*T*_ = *i*|**x**_*i*_) where **x**_*i*_ are the samples obtained for location *i* up until the response time, against the probability that they are caused by a distractor, *p*(*L*_*T*_ ≠ *i*|**x**_*i*_):

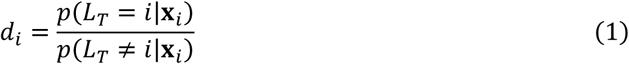

While this decision model is relatively simple, it provides a good intuition for observer behavior in the task (a more optimal model is provided in the Supplement but the conclusions do not depend on model choice). For this decision rule, the observer representation of distractor features learned from previous trials is related to response times:

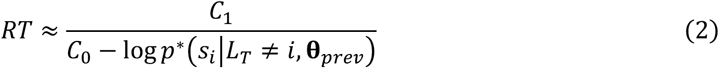

where *C*_0_and *C*_1_ are constants (see details in Methods). In words, there is an inverse relationship between response times and the approximate likelihood that a given stimulus is a distractor, *p*\*(*s*_*i*_|*L*_*T*_ ≠ *i*, **θ**_*prev*_), with the information obtained from previous trials described by a set of latent parameters, **θ**_*prev*_. When the probability that a stimulus at a given location (e.g., a test target) is a distractor is lower, response times are higher, and vice versa.

This model provides an important insight, namely, that observers’ representations are monotonically related to response times (Figure 2B). Hence, the relationship between the distribution parameters (mean, standard deviation, and skewness) reconstructed from RTs and from the true representation parameters would hold under any other monotonic transformation (for example, if RTs are log-transformed and the baseline is subtracted as we do in our analyses; see also Figure S1). In other words, response times can be used to approximately reconstruct observers’ representations of distractors and estimate their parameters.

**Figure 2.**
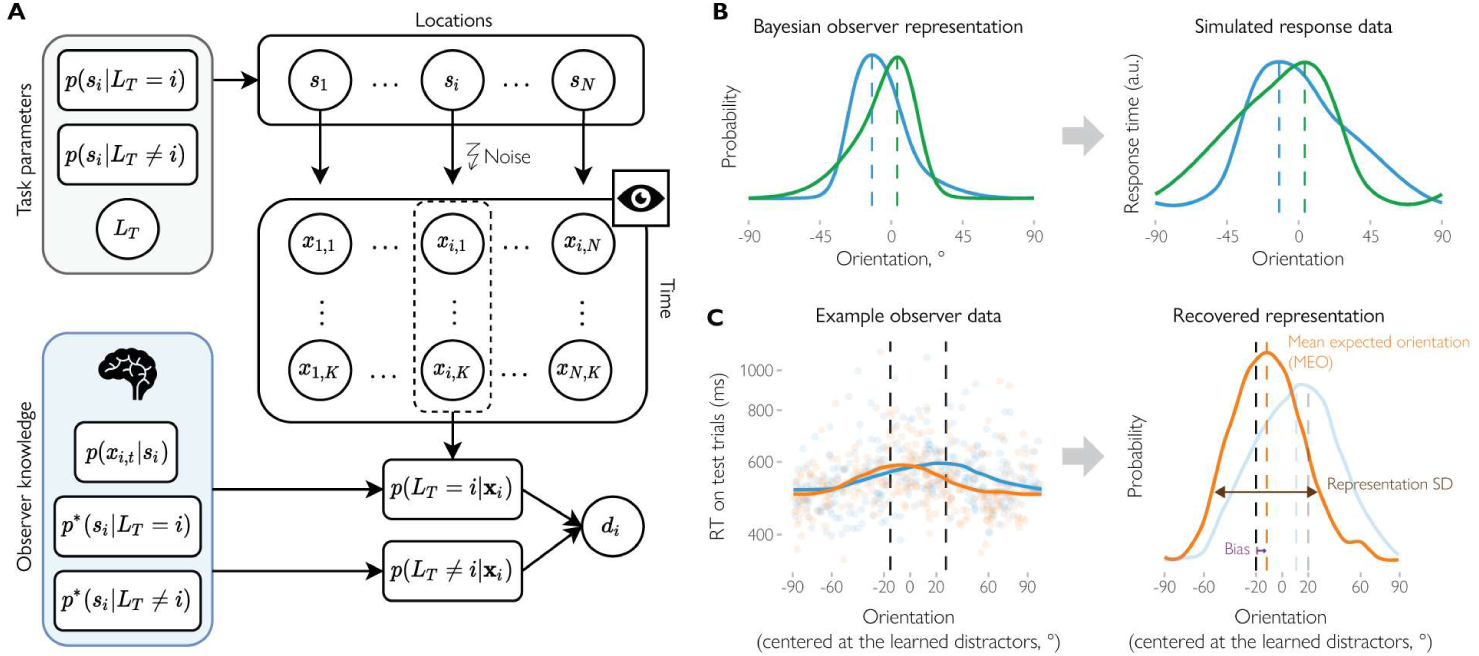
The Bayesian observer model provides a way of reconstructing distractor representations. **A**: The Bayesian observer model. The stimuli *s* … *s* at different locations are generated on each trial based on task parameters: the target feature distribution *p*(*s*_i_|*L*_*T*_ = *i*), the distractor feature distribution, *p*(*s*_i_|*L*_*T*_ ≠ *i*), and the target location *L*_*T*_. At each moment in time and for each location, observers obtain samples of sensory observations *x*_*i,t*_ corrupted by sensory noise, *p*(*x*_*i,t*_I*s*_*i*_). Using knowledge about the sensory noise distribution and the approximation of feature distributions for targets and distractors obtained during learning trials, *p*\*(*s*_i_|*L*_*T*_ = *i*) and *p*\*(*s*_i_|*L*_*T*_ ≠ *i*), observers compute probabilities that the sensory observations at a given location correspond to the target, *p*(*L*_*T*_ = *i*|**x**_*i*_), or a distractor, *p*(*L*_*T*_ ≠ *i*|**x**_*i*_). These probabilities are combined into a decision variable *d*_*i*_ used to make a decision or to continue gathering evidence if the currently available observations do not provide enough evidence for the decision (see details in Methods). **B**: The Bayesian observer model enables predictions about response times for a given representation of distractor stimuli (different example distributions are shown in blue and green). Crucially, there is a monotonic relationship between the two, with response times increasing with an increase in distractor probability. C: In our analyses, we used the monotonic relationship between probabilistic representations and response times to recover the representation of distractors (right) based on the response times on test trials (left). Here, the data from an example observer in the Spatial condition is split based on whether the target was located in the left (orange) or in the right (blue) hemifield. We then estimated the parameters of the representation, such as the mean expected orientation (dashed orange line), SD and across-distribution bias (the shift in the mean towards the other distribution relative to the true mean, shown by the dashed black line).

### Binding orientation probabilities to locations and colors

Having shown how observer response times should be related to the distractor representations, we now turn to the empirical data. By analyzing observers’ response times to different test targets, we were able to infer which orientations were most difficult to find, resulting in the longest response times. Crucially, we were able to reconstruct observers’ representations of the probability distributions that they were exposed to during learning trials (see Methods).

The experiments differed in the structure of the learning trials. There were three conditions in Experiment 1. The learning trials in the *Spatial* condition were organized so that distractor distributions in the left and the right hemifield differed to mimic the clustering of similar visual stimuli in the real world. In the *Color* condition, instead of spatial grouping, different distractor subsets were grouped by color while individual items were randomly distributed. Finally, in the *Baseline* condition items from the two distributions had the same color and were randomly distributed (Figure 1C).

Firstly, we report the results on the mean expected orientations (MEO) corresponding to the means of the recovered representations (Figure 2C). If observers ignore the separation of the two parts of the distribution, then MEO should match the mean of the overall distribution, but should differ between the distributions if the representations are bound to locations or colors. For example, if observers accurately learn the properties of the distributions, the MEO should be at +20° relative to the overall mean in the Spatial condition when the test line is presented in the hemifield that previously had distractors with an average relative orientation of +20°.

We found that in the Spatial condition, observers’ representations in each hemifield followed the actual physical distractor distribution. The estimated MEO relative to the overall mean was *M* = −14.02° (*SD* = 6.02) and *M* = 14.90° (*SD* = 5.14) for probes for clockwise (CW) and counterclockwise-shifted (CCW) distributions, respectively. The difference in MEO between the two distributions was much larger than zero (*b* = 28.94°, 95% HPDI = [25.34, 32.56], *BF* = 6.35 × 10^17^) showing that observers expected different orientations in different hemifields. We then computed the across-distribution bias by recoding the errors in MEO relative to the true mean for each distribution so that positive values correspond to shifts towards the other distribution. That is, the bias here represents by how much observers’ expectations deviated from the true mean orientation at a given location towards the mean orientation at the other location. For both hemifields there was a significant bias towards the other hemifield (*M* = 5.52°, 95% CI = [1.86, 9.14]). This shows that while observers represent the spatial separation between the two distributions, signals from the other hemifield still influence their responses.

But does spatial separation help observers to track the feature probabilities? In the Baseline conditions, locations of the CW and CCW distributions were chosen randomly for each learning trial. We repeated the analysis described above, comparing the response to test targets at the location that had CW and CCW orientations on immediately preceding trials. We expected to find stronger across-distribution biases as there was no separation between the distributions across trials. Importantly, the across-distribution bias was larger in the Baseline (bias *M* = 11.35°, 95% CI = [7.71, 15.00]) than the Spatial condition (effect of condition *M* = 5.84, 95% CI = [1.10, 10.58], *BF* = 108.24). In other words, the representations for each distribution were closer to the overall distribution mean in the Baseline than the Spatial condition. This argues that when the learned distributions are consistently presented at separate locations, observers can track them better than when they are mixed.

Do observers integrate information about orientation probabilities and color? In the Color condition, the locations of the test targets were counterbalanced with respect to their colors, so we should only find differences in MEO if observers formed an association between color and orientation. Indeed, we found that the MEOs for the two distributions differed (*b* = 7.35, 95% HPDI = [1.30, 13.06], *BF* = 148.04) although across-distribution biases were stronger (*M* = 16.30, 95% CI = [12.66, 19.86]) than in the Spatial condition (*M* = 10.78, 95% CI = [5.99, 15.54], *BF* = 6.56 × 10^4^). This means that if observers saw yellow lines shifted CW and blue lines shifted CCW relative to the overall distractor mean during learning trials, they learned this association which affected their response times on subsequent test trials. Importantly, this demonstrates that observers can integrate information about likely orientations with information about other features (in this case color), even if there is no spatial information to guide this integration.

### Encoding orientation probabilities at different spatial scales

Having established that observers associate information about most likely orientations with specific locations or colors, we then asked if we can uncover the origins of the observed biases by assessing the recovered representations in the Spatial condition in more detail (for this and later analyses, we increased the sensitivity of our analyses by combining the data from the Spatial group in Experiment 1 with an additional sample that performed the same task; see Methods). We computed MEO using the aggregated data from all participants for each location in the stimuli matrix in this condition. As Figure 3 shows, across-distribution biases were stronger closer to the boundary between the two hemifields. We then tested this observation by directly comparing MEOs for test trials with targets presented at the boundary (two central columns) between the hemifields against other test trials. We found that the bias was significantly larger at the boundary between the two distributions than in the other columns (*M* = 4.80° (*SD* = 6.99) and *M* = 9.04° (*SD* = 11.36), *b* = 4.23, 95% HPDI = [0.21, 8.32], *BF* = 42.34; Figure 3B). However, the biases were also significantly above zero outside the boundary (*BF* = 248). This suggests that the distribution representations are not homogenous and influence each other strongly when they are close in space, but this mutual influence also extends outside the immediate neighboring locations (see Discussion).

**Figure 3.**
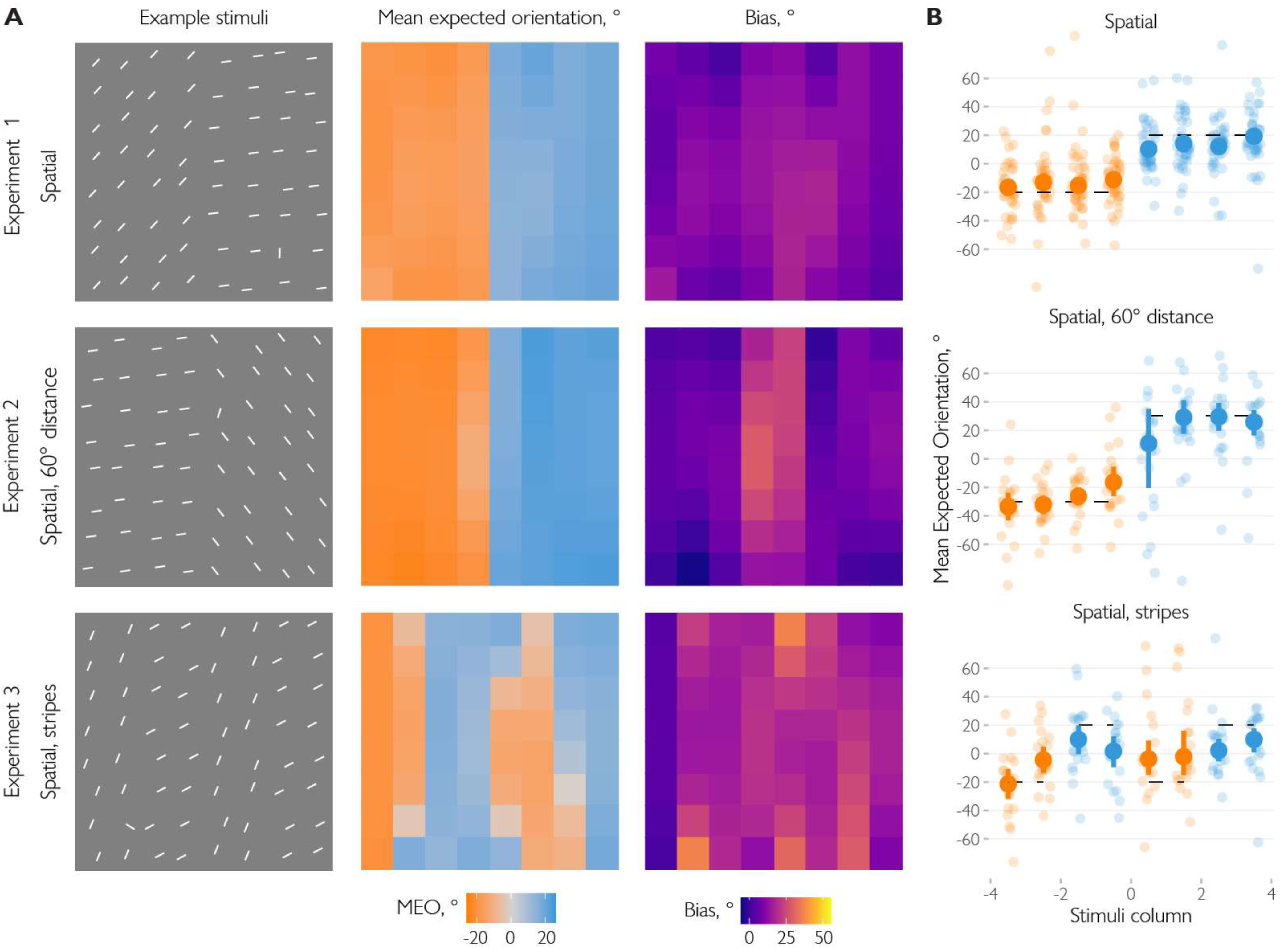
Spatial structure of probabilistic representations. **A**: Example stimuli (left column), recovered mean expected orientations (middle column) and the across-distribution biases in mean expected orientations relative to the true orientations at a given location (right column). The stimuli show a single learning trial from the search task in the corresponding experiment. The mean expected orientation (MEO) was then computed at each location relative to the overall average orientation in the preceding learning block. For presentation purposes, the data were rearranged so that the distribution in the left hemifield (or in the columns 1,2,5,6 in the stripes condition) was oriented clockwise relative to the overall mean. The biases in MEO were computed by subtracting the mean orientation for a given part of the distribution (e.g., at the left/right hemifield in the Spatial condition of Experiment 1) and recoding the resulting errors so that the positive values correspond to a bias towards the other distribution. **B**: Average MEO by column of stimuli matrix in the spatial conditions. Small dots show the data for individual observers, larger dots and bars show means and 95% CI, respectively. Dashed horizontal lines show the true means for a given part of the distribution.

### Bias strength depends on similarity and spatial arrangement

In two follow-up studies, we further investigated observers’ representations of spatially-grouped heterogeneous stimuli. In Experiment 2, we tested whether the similarity between the distributions along the tested feature dimension (orientation) affects the strength of the across-distribution biases. We hypothesized that the bias should be stronger when the stimuli from the two distributions are more likely to have the same cause in the external world. For example, the boundary effect in Experiment 1 might occur because the stimuli close in space are more likely to belong to the same object. By the same reasoning, if the two distributions are less similar, they are less likely to have the same cause, and the biases should be weaker.

To test this, we used the same spatial arrangement as in the Spatial condition in Experiment 1, but the distribution means were now 60° away from each other instead of 40° as in Experiment 1 (see example stimulus in Figure 3A). We found that again, MEO were close to their true values with *M* = 26.35° (*SD* = 13.43) and *M* = −27.65° (*SD* = 10.65) for distributions centred at 30° and −30° relative to the overall mean, respectively. Importantly, while there was a strong bias at the boundary between the distributions, *M* = 19.05° (*SD* = 27.27), *BF* = 8.36, it was absent at other positions (bias *M* = 0.60° (*SD* = 8.65), with *BF* = 4.12 in favor of no bias). Experiment 2, therefore, shows that reducing the similarity between the distributions eliminates the biases except for the immediately adjacent locations.

In Experiment 3, we tested whether an even more complex spatial arrangement would allow us to recover the “map” of observers’ expected orientations. To this end, the stimuli were organized in “stripes” of two matrix columns with two different distributions from Experiment 1 (with means separated by 40°) positioned at odd and even stripes (counterbalanced across blocks, Figure 3A). We found that observers expected clockwise-rotated orientations (*M* = 6.20°, *SD* = 9.91) at locations of stripes rotated 20° clockwise relative to the overall mean and counterclockwise-rotated orientations (*M* = −11.034°, *SD* = 17.11) at other stripe locations. However, the across-distribution bias (*M* = 11.70°, *SD* = 7.52) was stronger than in the Spatial condition in Experiment 1 (*b* = 5.90, 95% HPDI = [2.50, 9.33], *BF* = 4.30). This demonstrates that while separating distributions in space helps observers track distributions (as shown in Experiments 1 and 2), the effects of spatial organization decrease as the organization becomes more complex.

### Higher-order parameters of probabilistic representations

Next, we asked whether observers’ representations contain more information about the distributions than just their average? We used the reconstructed distractor representations (Figure 4A) to estimate their circular standard deviation and circular skewness. The former corresponds to the expected variability among distractors, while the latter quantifies their symmetry.

**Figure 4.**
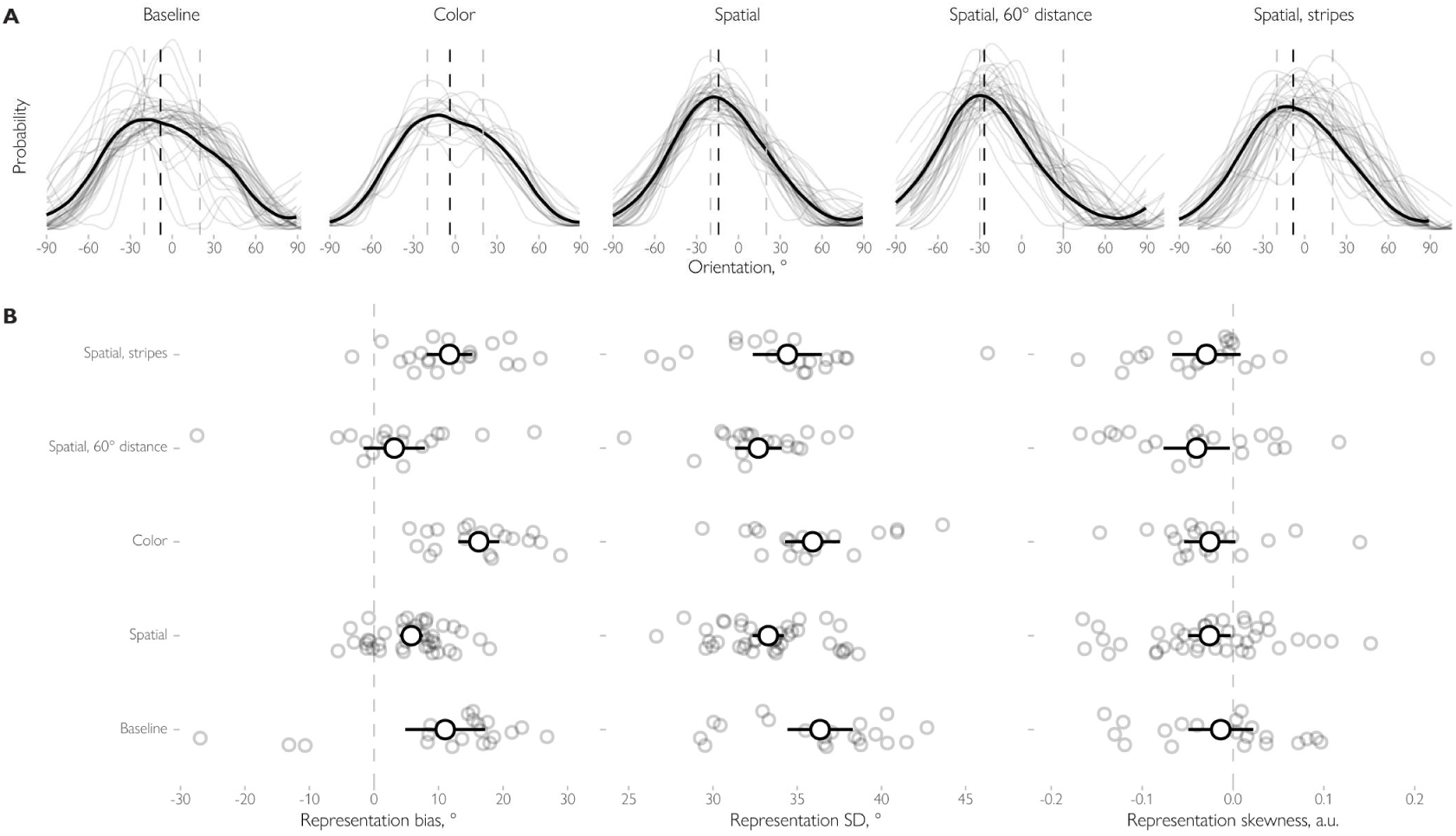
Recovered average representations and their parameters across experiments and conditions. **A**: The black curves show the average representation while representations for individual observers are shown in light gray. Dashed horizontal lines show the mean of the representation (black) and the true mean of the stimulus distributions (light gray). Note that the representations are aligned so that when two distributions are present, the true mean at the tested location is clockwise (−20° or −30°) while the other mean is counterclockwise (20° or 30° relative to the true mean). **B**: Estimated parameters (bias, SD and skewness). Large dots and errorbars show the mean across observers for a given parameter and the associated 95% confidence intervals. Smaller dots show data for individual subjects.

First, we hypothesized that if the variability of the distributions is encoded, then the expected variability would be higher when distractor distributions are less well separated. Indeed, we found that observers’ expectations about distractor variability differ between conditions (*BF* = 2.03 × 10^5^) with lower SD when the distractors were separated by hemifields (*M* = 33.3, 95% HPDI = [32.2, 34.4] for the Spatial condition with 40° separation and *M* = 32.7, 95% HPDI = [31.1, 34.2] for 60° separation) compared to other conditions (*M* = 35.9, 95% HPDI = [34.4, 37.5] in the color condition, *M* = 34.4, 95% HPDI = [32.9, 35.9] for the stripes arrangement condition). When the two distributions were less well separated, observers were more uncertain in their estimates, leading to distractor representations with higher SD’s (Figure 4B).

We also expected that the distribution presented at the tested location or in the tested color would weigh more highly in the resulting representation, causing an asymmetry. Alternatively, if observers only use the mean and variance to encode the distribution (as assumed by “summary statistics” accounts), then the represented distribution should be symmetric. We found that observers’ representations were asymmetric in all conditions, with a higher probability mass at the side corresponding to the distribution presented at the tested location or in the tested color, *M* = −0.03, 95% CI = [-0.04, −0.02]. Notably, however, no differences between conditions were found, *BF* = 1.99 × 10^−6^, indicating that symmetry is not affected by the way the distributions are organized in the display. In sum, observers represent not only the average stimulus values but also their variability, and the representations are skewed towards distributions presented at other locations or in different colors.

## Discussion

Our main hypothesis was that observers extract information about probabilities of visual features from heterogeneous stimuli and bind the resulting probabilistic representations with locations on the one hand and other features on the other. Our results support both these proposals very clearly, demonstrating how the visual system can build probabilistic representations of the visual world by extracting information about the features of complex heterogeneous stimuli.

A visual search task allowed us to uncover representations of heterogeneous distractors. We formulated a Bayesian observer model and demonstrated analytically and through simulations that response times are a monotonic function of observers’ expectations about distractor orientations, supporting earlier empirical findings (Chetverikov et al., 2016, 2017b, 2019, 2020). Using this knowledge, we were able to estimate the characteristics of observer representations – their means, precision, and skewness – and study how they vary depending on whether observers can associate them with locations or with other, task-irrelevant features, such as color.

We found that observers encode the feature distributions in scenes containing two different distributions. The representations generally follow the physical distribution of the stimuli for a given location or a given color, but importantly, observers are also biased towards the other distribution. The strength of the bias depends on the degree of separation between the distributions. When the distributions were separated in space, observers’ representations of one distribution were less influenced by the other distribution, compared to when they were separated by color or were intermixed (Baseline condition). Furthermore, as we found in Experiment 3, more complex spatial arrangements (“stripes”) increased the biases towards the other distribution. In sum, observers bind probabilistic representations of visual features to locations and other features, but such binding is not impenetrable, reminiscent of “illusory conjunctions” of discrete feature values (Treisman & Schmidt, 1982).

We were then able to recover the representation of the distribution at different spatial scales. We found that for spatial separation, the biases are stronger at the boundary between the two distributions. This is reminiscent of a hierarchical organization of information about feature probabilities within a scene proposed for perceptual ensembles (Alvarez, 2011; Haberman & Whitney, 2012). Such hierarchical ensemble models suggest that observers represent information about feature probabilities at different levels: for example, the orientation statistics at a particular location are combined to form a representation for a group of items, which are, in turn, combined to form an overall ensemble representation. Our results agree with this idea: the stimuli observers expect at a given location depend not only on what was previously shown at this location but also on stimuli presented at other locations. Crucially, biases were also present for the Color condition as well as for the non-boundary locations in the Spatial condition of Experiment 1. This indicates that the results cannot be explained by purely local summation of the inputs. It remains to be tested, whether there are actual separable representations of probability distributions at different levels, or just a unified spatio-featural map guiding observer responses.

We hypothesized that the representations should be more biased by each other when they are more likely to have the same cause in the external world. This could provide a normative explanation for the boundary effect: sensory input from adjacent locations is likely to be caused by the same object and should therefore be integrated while locations far away from each other should be treated separately. Similarly, for example, in multisensory integration studies, auditory and visual signals are less likely to be integrated when there is a large discrepancy in their locations (Körding et al., 2007; Shams & Beierholm, 2010). However, in Experiment 1 we found across-distribution biases at locations far from the other distribution. We reasoned that this is because the stimuli themselves are similar enough to be potentially caused by the same object, and the inputs are therefore integrated even from non-neighboring locations. In Experiment 2, we tested this explanation by asking if the similarity between the distributions themselves in the tested feature domain (orientation) also plays a role. We found that when the distributions were made more dissimilar, the biases were observed only at the boundary between the distributions but not at other locations.

That is, observers no longer take into account the input from non-neighboring locations, when stimuli are dissimilar. This supports the proposed normative explanation and suggests that the principles of information integration for heterogeneous visual inputs are the same as for other cases, such as multisensory integration or estimation of complex visual features (Landy et al., 2011; Shams & Beierholm, 2010).

We then tested if observers represent more than just the mean distractor orientation. We found that observers represent the distractor variability (i.e., the standard deviation or width of their representations), which varies in a predictable fashion with the separability between distractor distributions. When the distractor distributions are poorly separated (e.g., by color only or are organized in “stripes”), their representations are wider, indicating more uncertainty. Furthermore, the representations are asymmetric where the tail of the distribution corresponding to the orientations matching the tested location or color is fatter. That is, observers do not simply represent the distractors with a (biased) mean and variance, their representations have a complex shape with more relevant information (e.g., previous orientations at a tested location) weighted higher and less relevant information (e.g., previous orientations at the other locations) having lower weight, but still influencing the outcome.

These findings indicate that observers represent information about distractor features as a probability distribution rather than only in terms of the summary statistics, in contrast to popular ideas of simple “summary statistics”. For example, Treisman (2006) argued that statistical processing is a distinct mode of perceptual and attentional analysis of stimulus sets. She proposed that because of limited attentional capacity statistical summaries are generated that include the mean, variance, and perhaps the range. These summaries enable rapid assessment of the general properties and layout of natural scenes (Chong & Treisman, 2005; Emmanouil & Treisman, 2008). Similarly, Rahnev (2017; Yeon & Rahnev, 2020) argued that observers represent only a summary consisting of the most likely stimulus and the associated strength of evidence, and Cohen et al. (2016) used summary statistics to explain the richness of consciousness experience. Our results argue against such views, since the representations that are bound together are far more detailed than this implies. That is, the brain might instead approximate the visual input by using a complex set of parameters to provide accurate descriptions of feature probabilities (Freeman & Simoncelli, 2011; Rosenholtz, 2020).

A recent finding may explain why many previous studies have supported summary statistics proposals. Hansmann-Roth et al. (2021) reasoned that optimal behavior requires the encoding of full feature distributions, not only summaries, but observers might be unable to explicitly report the full distribution. This is analogous to how difficult it might be to verbally describe the variety of colors of an apple without resorting to simplifications (see Figure 1A). Hansmann-Roth et al. tested observers’ representations both implicitly and explicitly and while explicit judgments were limited to the mean and variance of feature distributions, implicit measures revealed detailed representations of the same distributions. More information was therefore available to observers than studies of summary statistics, that have mostly relied on explicit measures, have indicated. Crucially, Hansmann-Roth et al. were able to uncover why this is: revealing these detailed representations requires implicit methods, such as we use here.

In our experiments, observers learn the distractor feature by combining inputs from heterogeneous stimuli across several trials in each block, and it can be argued that this is different from perceiving a single stimulus on a single trial. However, the visual cortex aggregates information at many different timescales (de Lange et al., 2018). Even on a single trial, perception unfolds in time and at each moment is dependent on what has been seen before. And even for a simple stimulus, the visual cortex receives inputs from many retinal neurons that are affected by processing noise, potentially indistinguishable from the input from varying features. Indeed, this is why stimulus variability (“external noise”) is often used to manipulate visual uncertainty (Barthelmé & Mamassian, 2009; Hénaff et al., 2020). We therefore believe that distinguishing “simple” and “complex” perception is impossible. However, our results clearly show that information about feature probabilities is available for visually-guided behavior.

Taken together, our results show that observers can not only encode probabilities of features from heterogeneous stimuli in detail but also integrate them with both locations and other features that have different distributions. These results arguably represent the strongest support yet for the long-standing idea that the brain builds probabilistic models of the world (Chetverikov et al., 2017a; Fiser et al., 2010; Knill & Pouget, 2004; Orhan & Ma, 2015; Rao et al., 2002; Sahani & Dayan, 2003; Tanrıkulu et al., 2021) and show that probabilistic representations can serve as building blocks for object and scene processing. Notably, such representations are not simply limited to summary statistics (e.g., a combination of mean and variance (Cohen et al., 2016)). Our results also indicate that observers do not represent physical stimuli precisely, but instead construct an approximation influenced by input from other stimuli. This probabilistic perspective stands in sharp contrast to views where discrete features of individual stimuli are *either* bound together to form objects or processed “statistically” (Rosenholtz, 2020; Treisman, 2006). Instead, we suggest that the probabilistic representations are automatically bound to locations and other features since such binding occurred even though it was not required in the task. Probabilistic representations are therefore not acquired in isolation but constitute an integral part of perception.

## Methods

### Participants

In total, eighty observers (fifty female, age *M* = 23.10) participated in the experiments. Twenty observers (ten female, age *M* = 25.45) participated in the first experiment (Baseline, Spatial, and Color conditions) split across two sessions. Twenty observers (fourteen female, age *M* = 25.00) participated in Experiment 2 (“Spatial, 60° distance”) and another twenty (thirteen female, age *M* = 25.45) in Experiment 3 (“Spatial, stripes”). Finally, the data from additional twenty observers (thirteen female, age *M* = 16.50) were collected for the Spatial condition of Experiment 1 to increase the sensitivity of the spatial analyses.

All were staff or students at the Faculty of Psychology, St. Petersburg State University, Russia, or the University of Iceland, Iceland. The experiment was approved by local ethics boards and was run in accordance with the Helsinki declaration. Participants at St. Petersburg State University were rewarded with 500 rubles (approx. 8 USD) per hour each, participants at the University of Iceland participated without additional reward. All gave their informed consent before participating. The participants were naïve to the purposes of the studies. Participants were given ample time for training until they felt comfortable doing the task (the training time ranged from 5 minutes to one hour depending on the participant).

### Procedure

In *Experiment 1*, each participant performed a search task in five conditions. In each condition on each trial, observers were presented with 8×8 matrices of 64 lines (line length: 0.71° of visual angle; matrix size: 16×16°; uniform noise of ±0.5° was added to each line coordinate). The goal was to find the odd-one-out line whose orientation differed most from the others. Sessions were separated into blocks of 5 to 7 learning trials followed by 1 or 2 test trials (the number of trials chosen randomly for each block; the variation in the number of trials was introduced to decrease the effect of temporal expectations(Shurygina et al., 2019)). During learning trials, the overall mean of distracting items varied randomly with half of the distractors drawn from one distribution and the other half from another distribution with the properties of distributions differing between conditions:

*Baseline*: two truncated Gaussian distributions with SD = 10° and range of 40°, with means separated by 40° (±20° relative to the overall mean), all stimuli had the same color (white), positions for each line within the matrix were chosen randomly.

Spatial: two distributions (either a truncated Gaussian with SD = 10° and a range of 40° or uniform with the range of 40°) with means separated by 40° (±20° relative to the overall mean), all stimuli had the same color (white), one distribution was shown in the left half of the matrix, the other in the right half.

Color: the same distributions as in the Spatial condition were used, but lines drawn from one distribution were blue, while lines from the other distribution were yellow. Positions for each line within the stimuli matrix were chosen randomly.

In all cases, two lines were added to each distractor distribution with their orientation equal to the minimal and maximal values from that distribution range. As a result, Gaussian and uniform distributions always had the same range. Target orientation on each trial was drawn randomly from a uniform distribution ranging between 60° and 120° relative to the mean distractor orientation.

On test trials, distractors came from a single Gaussian distribution with SD = 10° (range-restricted in the same way as described above), while target orientation was determined in the same way as on the prime trials. In the color condition, half of the lines from that distribution were blue, half were yellow.

The Baseline condition had 2304 trials, while the Spatial and Color conditions had 5376 trials each with the higher number of trials used in the latter case to counterbalance additional factors (distribution type combinations).

*Experiments 2 and 3* generally followed the same procedure as the Spatial condition of Experiment 1. In Experiment 2 the means of the distributions were separated by 60° (±30° relative to the overall mean) instead of 40° in Experiment 1. In Experiment 3, the two distributions were separated by 40°, as in Experiment 1, but arranged in “stripes” so that the lines drawn from the first distribution were positioned in the 1^st^, 2^nd^, 5^th^, and 6^th^ columns of the stimuli matrix while the other columns were populated with lines from the second distribution.

### Data processing

For our main analyses of interest, incorrect responses were excluded and response times were log-transformed and centered by subtracting the mean for each participant. Then, to reduce the noise in RT measurements, spatial and featural confounders were removed. First, the effect of the distance between target locations on consecutive trials and the effect of the target location were removed by regressing out the fifth-degree polynomials of the absolute distance (in degrees of visual angle) between the target locations on the current and the previous trials and the current targets horizontal and vertical coordinates. Then, we also removed potential influences from the well-known oblique effect (the search speed differs between oblique and cardinal stimuli (Chetverikov et al., 2017a; Wolfe et al., 1999) by regressing out the fifth-degree polynomials of target and distractor obliqueness computed as an absolute distance in degrees to the nearest cardinal orientation. The regression was run separately for each experiment and condition.

To reconstruct observers’ distractor representations, we used the response times on the first test trial in each block. We then converted response times as a function of the similarity between the test target and previous distractor mean to a probabilistic representation and estimated its parameters.

To convert the noisy response times into probabilities, we first smoothed RT as a function of the test target and previous distractor mean using the local regression approach (a generalization of the moving average) for each observer in each condition. To account for circularity, we appended 1/6 of the data from each end of the orientation space to the opposite end before smoothing. In analyses applied to each stimulus location, we further assumed that RTs are a smooth function of the stimuli matrix row within the local regression while columns of the stimuli matrix were treated independently. We then transformed a smoothed RT function into a probability mass function by subtracting the baseline and normalizing to one. Finally, we computed the parameters of the recovered probabilistic representation: the mean expected orientation (circular mean), circular standard deviation, and circular skewness as defined by Pewsey(2004). Note that under the hypothesized Bayesian observer model, the estimated standard deviation and skewness are monotonically related to the true parameters of the distractor representation but are not identical to it (additionally confirmed in simulations, Figure S1).

### Data analysis

Unless stated otherwise, we used Bayesian hierarchical regression with *brms* (Bürkner, 2017) package in R. Note that while we include Bayes factor values in the description of the results, we were mostly interested in measuring the effects of the variables of interest in our models, hence the models included the default flat (uniform) priors for regression coefficients. Given that Bayes factors are heavily prior-dependent, we believe that the information provided by the 95% highest-density posterior intervals (HDPI) is more useful for judging the results than the Bayes factors. To make sure that the conclusions are not dependent on the particular analytic approach, we repeated the analyses using the conventional frequentist statistical test with the same results (the report using this approach is provided alongside the data in an online repository, see *Data availability* statement).

### Bayesian observer model

In our experiments, participants located a target among a set of distractors and indicated if it is in the top or the lower part of the stimuli matrix. On each trial, the experimenter sets the task parameters, namely, parameters of the target distribution, *p*(*s*_*i*_|*L*_*T*_ = *i*), and parameters of the distractor distribution, *p*(*s*_*i*_|*L*_*T*_ ≠ *i*), for each location *i* = 1 … *N* in the stimuli matrix as well as the target location, *L*_*T*_. These parameters were then used to generate the stimuli at each location, *s*_*i*_.

Neither the task parameters nor the stimuli are known to the Bayesian observer. Instead, at each moment in time *t*, the observer obtains sensory observations at each location, *x*_*i,t*_. These observations are not identical to the stimuli because of the presence of sensory noise, *p*(*x*_*i,t*_|*s*_*i*_). That is, a given stimulus might result in different sensory responses, and, conversely, a given sensory observation might correspond to different stimuli. We assume that the observations are distributed independently at each location and at each moment in time.

To make an optimal decision in a particular task, the observer needs to know the relationship between the sensory observations and the task-relevant quantities. For the visual search task used in our study, we assumed that observers compare for each location the probability that the sensory observations are caused by a target present at that location, *p*(*L*_*T*_ = *i*|**x**_*i*_) where **x**_*i*_ = {*x*_*i*,_, *x*_*i*,2_, …, *x*_*i,t*=*K*_} are the samples obtained for location *i* up until the time *K*, against the probability that they are caused by a distractor, *p*(*L*_*T*_ ≠ *i*|**x**_*i*_):

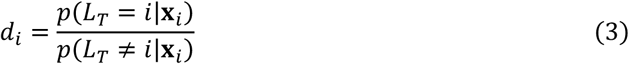

The observer then decides that a given item is a target as soon as the decision variable at a given location reaches a certain threshold *B*. Although this decision rule is not fully optimal, because the observer makes a decision for each item individually, it greatly reduces the task complexity, and we believe that it allows for a more realistic model (the simulations based on a more complex but more optimal model are described in the supplement and lead to identical conclusions).

The observer can compute the probability of hypotheses *L*_*T*_ = *i* and *L*_*T*_ ≠ *i* given the sensory data using the Bayes rule:

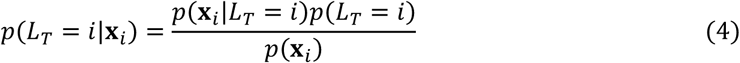

In words, the probability of a hypothesis that a target is at the given location, *L*_*T*_ = *i*, for a set of sensory observations **x**_*i*_ is equal to the likelihood of the data given this hypothesis multiplied by a prior probability for this hypothesis *p*(*L*_*T*_ = *i*) and divided by the probability of the observations *p*(**x**_*i*_).

Assuming that the prior 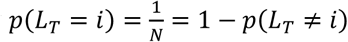 is the same for all locations, the decision variable can then be rewritten in log-space as the difference in the log-likelihoods in favor of the two hypotheses:

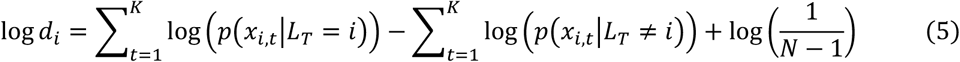

What are the probabilities of sensory observations under each hypothesis, *p*(*x*_*i,t*_|*L*_*T*_ = *i*) and *p*(*x*_*i,t*_|*L*_*T*_ ≠ *i*)? To compute them, the observer needs to take into account how the stimuli are distributed under each hypothesis and how the sensory noise is distributed for each stimulus. We assume that the sensory noise distribution is known for the observer through long-time exposure to the visual environment (that is, the observer knows *p*(*x*_*i,t*_|*s*_*i*_)).

However, to determine how probable it is that sensory observations correspond to the search target, the observer must also know what defines targets and distractors. The experimenter knows that only certain orientations describe a target, but the observer is not omniscient and does not know the true distributions of target and distractor stimuli, approximating them instead as *p*\*(*s*_*i*_|*L*_*T*_ = *i*) and *p*\*(*s*_*i*_|*L*_*T*_ ≠ *i*). Then the probability of sensory observations under each hypothesis can be computed as:

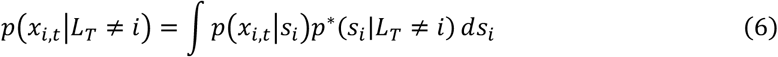

The probability distributions *p*\*(*s*_*i*_|*L*_*T*_ = *i*) and *p*\*(*s*_*i*_|*L*_*T*_ ≠ *i*) correspond to the observer’s approximate representation of target and distractor distributions. Notably, each of them can be further separated into the representation based on the previous trials and the one based on the current trial:

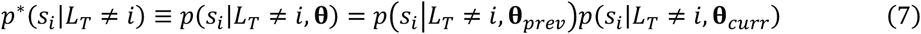

with **θ** = {**θ**_*prev*_, **θ**_*curr*_} corresponding to the independent latent variables describing the parameters of the previous and the current trial by the observer (similar equations related to targets are omitted for brevity). In our experiments, by design, the parameters of the current trial are controlled with respect to the current stimuli (i.e., the distractors on the current test trial are drawn from a distribution with a mean from 60° to 120° off the current test target). Hence, only *p*(*s*_*i*_|*L*_*T*_ ≠ *i*, **θ**_*prev*_) matters for relative changes in response times.

In our analyses, we wanted to reconstruct the representation of distractor stimuli using the response times for different test targets. Because the decision time is proportionate to the number of samples when the sampling frequency is constant, we aimed to relate the number of samples *K* to an observer’s approximate representation of distractors based on the previous trials *p*(*s*_*i*_|*L*_*T*_ ≠ *i*, **θ**_*prev*_).

Assuming that the sensory observations are obtained with high frequency, we can approximate the total evidence in favor of a given hypothesis:

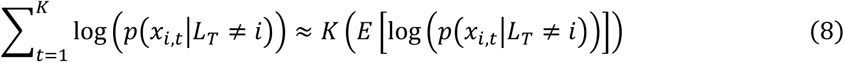

We expect sensory noise to be low compared to the noise in the target and distractor representations. Then, the following approximation is valid:

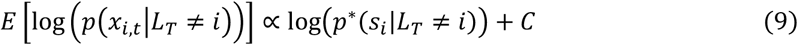

where *C* is a constant. Similar derivations can be used for the total evidence for the alternative hypothesis *p*(*x*_*i,t*_|*L*_*T*_ = *i*).

Then, given that a decision is made when log *d*_*i*_ = log *B*:

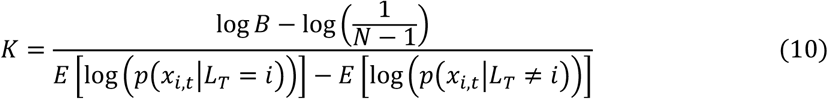

Given that the target and distractor parameters are independently manipulated in the experiment, *E* [log (*p*(*x*_*i,t*_|*L*_*T*_ = *i*))] can be treated as a constant. Similarly, *p*(*s*_*i*_|*L*_*T*_ = *i*, **θ**_*curr*_) would be constant as discussed above. Given that *RT* ∝ *K*, we can then approximate is as follows:

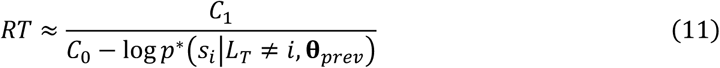

and

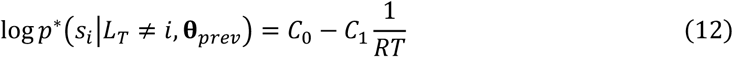

where *C*_0_ and *C*_1_ are constants. In words, there is an inverse linear relationship between the likelihood that a given stimulus is a distractor (in log-space) and the response times. When this likelihood increases, response times decrease.

This model provides an important insight, namely, that observers’ representations are monotonically related to response times. Hence, even though *C*_0_ and *C*_1_ are unknown, the relationship between the moments (mean, standard deviation, and skewness) of observers’ representations reconstructed from RT and the true representations would hold under any other monotonic transformation (for example, RTs are log-transformed and the baseline RTs are subtracted as we do in our analyses).

## Supporting information

in the supplement

## Acknowledgments

Supported by a grant from the Icelandic Research Fund (IRF #173947-052) and a grant from the Russian Foundation for Basic Research (RFBR, #15-36-01358). AC was supported by the Radboud Excellence Initiative. We are grateful to Alena Begler for the help with data collection and to James Cooke for his invaluable comments on the models included in the manuscript.

## Data availability

The data and scripts used for the data analysis in this paper are available from https://osf.io/5pfyn/.

## Notes

https://osf.io/5pfyn/

### Competing Interest Statement

The authors have declared no competing interest.

### Summary of Updates

The title and the abstract were updated for clarity.

