## Supplementary material for "Probabilistic representations as building blocks for higher-level vision": in the supplement

### Supplement 1. Bayesian observer model combining information across locations.

The model reported in the main text presents a simplified version of the decision-making process assuming that stimuli at each location are analyzed separately. We believe that such a model might be more realistic as it greatly simplifies the computations that observers have to make. However, for the sake of completeness, here we briefly describe a more complex conditionally-optimal memory-guided Bayesian observer model. We refer to this model as conditionally optimal for two reasons. First, a memory-guided observer is by definition not fully optimal in our task, where the test trial parameters are unrelated to the previous learning trials. However, given that the task parameters repeat throughout learning trials, using the information from the previous trials might be beneficial when the observer does not know that the trial parameters have changed. Secondly, we assume that the observer's learning or memory about the stimuli features might not be ideal, hence they use the approximations of feature distributions. We show that under this more complex and more optimal model, the predictions with respect to the monotonic relationship between the response times and expected distractor probabilities stay the same.

***Task structure.*** Participants have to locate a target among a set of distractors and indicate if it is in the top or in the lower part of the stimuli matrix. The experimenter sets the task parameters for each trial, namely, the target distribution, $p(s_i|L_T = i)$, and the distractor distribution, $p(s_i|L_T \neq i)$, for each location $i = 1 \dots N$ in the stimuli matrix (with top half having indices from 1 to $N/2$ and the bottom half from $\frac{N}{2} + 1$ to $N$) as well as the target location ($L_T$), to generate the stimuli ($s_i$) at each location. Here, $L_T = i$ and $L_T \neq i$ indicate that the target is or is not at location *i*, or in other words, that the target location is or is not *i*, respectively.

*Ideal observer model.* At each moment in time $t = 1 \dots K$ (with *K* as the decision moment) and at each location *i*, the observer obtains sensory observations $x_{i,t}$ corrupted by the presence of sensory noise:

$$p(x_{i,t}|s_i) = f_{VM}(x_i; s_i, \kappa_s)$$

where $f_{VM}$ is a von Mises distribution density with concentration parameter $\kappa_s$ quantifying the amount of noise. We assume that the observations are distributed independently at each location and at each moment in time:

$$p(\mathbf{X}|\mathbf{s}) = \prod_{i=1}^{N} p(\mathbf{x_i}|s_i) = \prod_{i=1}^{N} \prod_{t=1}^{K} p(x_{i,t}|s_i) \quad \text{(S13)}$$

To make an optimal decision in a particular task, the observer needs to compare the probability that a target is located in the upper half of the stimuli matrix with a probability that it is located in the lower half:

$$d = \frac{p(C = 1|\mathbf{X})}{p(C = 2|\mathbf{X})} \quad \text{(S14)}$$

where $C = 1$ and $C = 2$ correspond to the two hypotheses about the target location. After applying the log transformation, the decision variable can be expressed as a difference in the amount of evidence for the two hypotheses:

$$\log d = \log p(C = 1|\mathbf{X}) - \log p(C = 2|\mathbf{X}) \quad \text{(S15)}$$

The decision time assuming a certain threshold $B$ can then be found as a time $K$ when the decision variable reaches the threshold. The average decision time can be found by estimating when the expectation of $\log d$ becomes equal to $\log B$:

$$K = \frac{\log B}{E[\log p(C = 1|\mathbf{X})] - E[\log p(C = 2|\mathbf{X})]} \tag{S16}$$

The probabilities for each hypothesis $C = 1$ and $C = 2$ can be found using the Bayes rule. For example, for $C = 1$:

$$p(C = 1|\mathbf{X}) = \frac{p(\mathbf{X}|C = 1)p(C = 1)}{p(\mathbf{X})} \tag{S17}$$

Because the observer does not know what stimuli are presented and only knows the sensory observations, the likelihood $p(\mathbf{x}|C = 1)$ needs to be computed by averaging (marginalizing) over the unknown stimuli values:

$$p(\mathbf{X}|C = 1) = \int p(\mathbf{X}|\mathbf{s})p(\mathbf{s}|C = 1)d\mathbf{s} \tag{S18}$$

Because the target can be only present at one location, the likelihood $p(\mathbf{x}|C = 1)$ is computed by summing over the possibilities of finding a target at each particular location:

$$p(\mathbf{X}|C = 1) = \sum_{i=1}^{\frac{N}{2}} \int p(\mathbf{X}|\mathbf{s})p^*(\mathbf{s}|L_T = i, \boldsymbol{\theta})d\mathbf{s} \tag{S19}$$

where similarly to the main text, we use an asterisk to denote probability distributions as approximated by the observer through a set of parameters related to previous and current trials $\boldsymbol{\theta} = \{\boldsymbol{\theta}_{prev}, \boldsymbol{\theta}_{curr}\}$. That is, we assume that the observer is unaware of the true distributions $p(s_i|L_T = i)$ and $p(s_i|L_T \neq i)$ and approximates them instead using the information available.

If a target is at location *i,* it cannot be anywhere else. Hence:

$$p^*(\mathbf{s}|L_T = i, \boldsymbol{\theta}) = p^*(s_i|L_T = i, \boldsymbol{\theta}) \prod_{j \neq i}^{N} p^*(s_j|L_T \neq j, \boldsymbol{\theta}) \tag{S20}$$

Using Eq. S20, it can be further shown that:

$$\int p(\mathbf{X}|\mathbf{s})p^*(\mathbf{s}|L_T = i, \boldsymbol{\theta})d\mathbf{s} = \left[\prod_{j}^{N} \int p(\mathbf{x}_\mathrm{j}|s_j)p^*(s_j|L_T \neq j, \boldsymbol{\theta})ds_j\right] \frac{\int p(\mathbf{x}_\mathrm{i}|s_i)p^*(s_i|L_T = i, \boldsymbol{\theta})ds_\mathrm{i}}{\int p(\mathbf{x}_\mathrm{i}|s_i)p^*(s_i|L_T \neq i, \boldsymbol{\theta})ds_\mathrm{i}} \tag{S21}$$

Note that the product in the square brackets is the same for all locations, and the remaining part of the equation is a ratio of the probability that the measurements at a given location are from the target against the probability that they are from the distractor, similarly to the model described in the main text.

The probability that a given stimulus is a target (or a distractor) depends on both the previous and the current trial:

$$p^*(s_i|L_T = i, \boldsymbol{\theta}) = p^*(s_i|L_T = i, \boldsymbol{\theta}_{prev})p^*(s_i|L_T = i, \boldsymbol{\theta}_{curr}) \tag{S22}$$

For each location and each location-specific hypothesis $L_T = i$ and $L_T \neq i$, the current trial parameters need to be computed separately because of the nature of the odd-one-out task. A target is defined as the item most different from the distractors. For simplicity, we assumed that observers use the following circular normal approximation for the distractors at the current trial based on the sensory observations:

$$p^*(s_i|L_T \neq i, \boldsymbol{\theta}_{curr}) = f_{VM}(s_i; \hat{\mu}_{j\neq i}, \hat{\kappa}_{j\neq i}) \quad \text{(S23)}$$

In words, when the observer needs to estimate, how likely it is that the stimulus at location *i* is a distractor, the observer approximates the distribution of stimuli as a von Mises (circular normal) distribution based on the sensory observations from other locations.

The observer might use the knowledge that the target distribution in the task design is on average 90° away from the mean of distractors. We again assume a von Mises approximation:

$$p^*(s_i|L_T = i, \boldsymbol{\theta}_{curr}) = f_{VM}(s_i; \hat{\mu}_{j\neq i} + 90°, \kappa_T) \quad \text{(S24)}$$

where $\kappa_T$ is the expected precision of the target distribution. In contrast to the distractor distribution precision that could be guessed based on the samples on the current trial ($\hat{\kappa}_{j\neq i}$), the target distribution precision cannot be estimated on a single trial (there is only one target stimulus in a given trial) and has to be based on the other sources of information (e.g., learning throughout the experiment).

Given that the measurement noise is independent across locations, the likelihood of the hypothesis *C* = 1 can be further expressed as:

$$p(\mathbf{X}|C=1) = \left[\prod_{j=1}^{N} \int (\mathbf{x}_j|s_j) p^*(s_j|L_T \neq j, \boldsymbol{\theta}) ds_j\right] \sum_{i=1}^{\frac{N}{2}} \frac{\int p(\mathbf{x}_i|s_i) p^*(s_i|L_T = i, \boldsymbol{\theta}) ds_i}{\int p(\mathbf{x}_i|s_i) p^*(s_i|L_T \neq i, \boldsymbol{\theta}) ds_i} \quad \text{(S25)}$$

Then, assuming that the prior probability of each decision alternative is the same, the decision variable can be expressed in log-space as:

$$\log d = \log\left(\sum_{i=1}^{\frac{N}{2}} \frac{\int p(\mathbf{x}_i|s_i) p^*(s_i|L_T = i, \boldsymbol{\theta}) ds_i}{\int p(\mathbf{x}_i|s_i) p^*(s_i|L_T \neq i, \boldsymbol{\theta}) ds_i}\right) - \log\left(\sum_{i=\frac{N}{2}+1}^{N} \frac{\int p(\mathbf{x}_i|s_i) p^*(s_i|L_T = i, \boldsymbol{\theta}) ds_i}{\int p(\mathbf{x}_i|s_i) p^*(s_i|L_T \neq i, \boldsymbol{\theta}) ds_i}\right) \text{(S26)}$$

The decision time assuming a certain threshold *B* can then be found as a time *K* when the decision variable reaches the threshold.

**Simulations.** To estimate the behavior of the observer using this model, we simulated the decision-making process and estimated the mean response times while varying the properties of the distractor representation $p^*(s_i|L_T \neq i, \boldsymbol{\theta}_{prev})$. The task parameters were based on the actual experiment design. We used 36 stimuli for each trial with one stimulus being the test target ($s_{L_T}$) and the rest being the distractors. The distractors on each simulated trial were distributed as $p(s_i|L_T \neq i) = f_{VM}(s_i; \mu_D, \kappa_D)$ where $\mu_D \sim U(s_{L_T} + 60°; s_{L_T} + 120°)$ (that is, the mean of distractors is set to 60° to 120° away from the test stimulus) and $\kappa_D = 8.7$ (approximately equivalent to the standard deviation of 10° in orientation space). The sensory observations were assumed to be noisy ($\kappa_S = 2$, approximately equivalent to the standard deviation of 24° in orientation space; note that this is the noise level for samples collected at each moment in time). The observers' target representation was assumed to be linked with to the distractor representation as $p^*(s_i|L_T = i, \boldsymbol{\theta}_{prev}) = f_{VM}\left(s_i; \mu_{D_{prev}}, \kappa_T\right)$ with $\kappa_T = 3.35$ (based on a normal approximation to a uniform target distribution with 60° range used in the experiments). The same $\kappa_T$ was used for target-related computations based on the current trial data (Eq. S24). The decision threshold was set to $\log B = 4.60$ assuming a 1% probability of error if the observer assumptions are correct. For each test target from 1° to 180° in half-degree steps, we simulated 56 trials for each combination of distractor representation parameters.

We ran simulations for the wrapped skewed normal distribution with the mean varied from -60° to 60° in 20° steps, while the standard deviation varied from 20° to 60° in 10° steps, and skew varied from -10 to 10 in steps of 2. The results of the simulations (Figure S2) confirmed the findings obtained with a simplified model: the means are recovered precisely while for standard deviation and skewness the monotonic relationship holds.

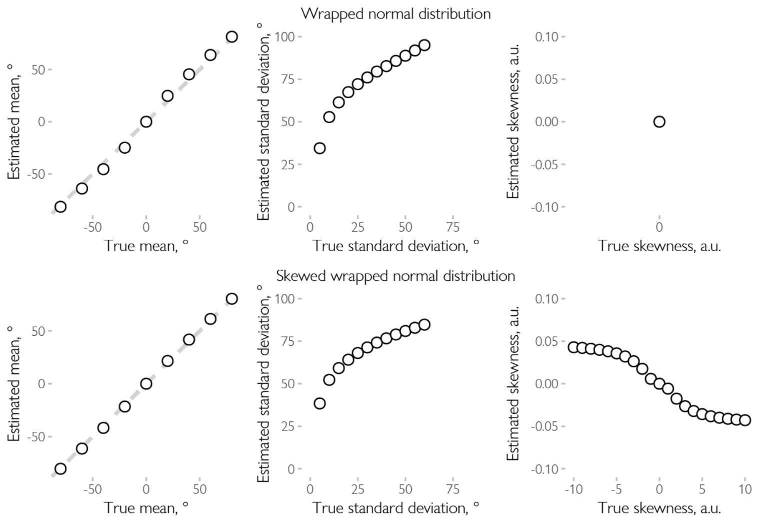


Figure S1. Simulated parameters under the simplified Bayesian observer model. We simulated the response times under the assumptions of the simplified Bayesian observer model described in the main text and applied the same approach as used for the real data to see if the assumed monotonic relationship between the true parameters and the recovered parameters holds. Firstly, we used a simple wrapped normal (top) with means varying from -80° to 80° in 20° steps and standard deviation from 5° to 60° in 5° steps. For each parameter combination the RT were computed using Eq. 2. We then estimated the parameters of the recovered distribution. As is evident from the plots, the mean estimates were identical to the true mean while the standard deviation was overestimated but the overall monotonic relationship held. The skewness estimate was at zero as expected for the symmetric wrapped normal distribution. Secondly, we simulated the data using the skewed normal distribution (Pewsey, 2008) with means again varying from -80° to 80° in 20° steps, scale parameter varying from 5° to 60° in 5° steps, and skewness parameter varying from -10 to 10 in steps of 1. For the means and standard deviations, the conclusions were the same as for the wrapped normal distribution. Similarly, skewness estimates followed monotonically the changes in the true skewness parameter (note that the sign of the estimated circular skewness is the opposite of the skewness parameter of the skewed wrapped normal distribution because of how it is defined, see Pewsey, 2004). In sum, the mean estimates match the true means, and the standard deviation and skewness estimates monotonically depend on the true standard deviation and the skewness parameters.

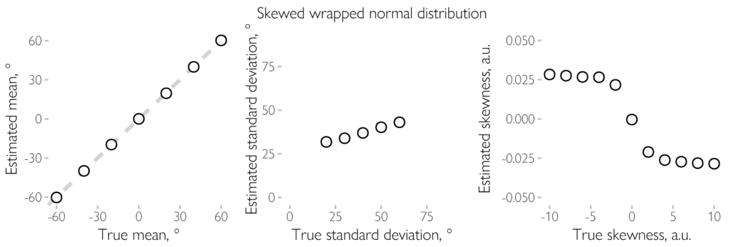


Figure S2. Simulated parameters under the more optimal Bayesian observer model. We simulated the response times under the assumptions of the more complex Bayesian observer model described in the Supplement applied the same approach as used for the real data to see if the assumed monotonic relationship between the true parameters and the recovered parameters holds. The results were similar to the simulations with the simplified model (Figure S1). The mean estimates were identical to the true mean, while for the standard deviation and skewness the monotonic relation holds (note that the sign of the estimated circular skewness is the opposite of the skewness parameter of the skewed wrapped normal distribution because of how it is defined, see Pewsey, 2004). In sum, the mean estimates match the true means, and the standard deviation and skewness estimates monotonically depend on the true standard deviation and the skewness parameters.
